# The landscape of *SETBP1* gene expression and transcription factor activity across human tissues

**DOI:** 10.1101/2023.08.08.551337

**Authors:** Jordan H. Whitlock, Elizabeth J. Wilk, Timothy C. Howton, Amanda D. Clark, Brittany N. Lasseigne

## Abstract

**Background:** The SET binding protein 1 (*SETBP1*) gene encodes a transcription factor (TF) involved in various cellular processes. Distinct *SETBP1* variants have been linked to three different diseases. Germline variants cause the ultra-rare pediatric Schinzel Giedion Syndrome (SGS) and *SETBP1* haploinsufficiency disorder (*SETBP1*-HD), characterized by severe multisystemic abnormalities with neurodegeneration or a less severe brain phenotype accompanied by hypotonia and strabismus, respectively. Somatic variants in *SETBP1* are associated with hematological malignancies and cancer development in other tissues in adults.

**Results:** To better understand the tissue-specific mechanisms involving *SETBP1*, we analyzed publicly available RNA-sequencing data from the Genotype-Tissue Expression (GTEx) project. We found *SETBP1*, and its known target genes were widely expressed across 31 adult human tissues. K-means clustering identified three distinct expression patterns of SETBP1 targets across tissues. Functional enrichment analysis (FEA) of each cluster revealed gene sets related to transcription regulation, DNA binding, and mitochondrial function. TF activity analysis of SETBP1 and its target TFs revealed tissue-specific TF activity, underscoring the role of tissue context-driven regulation and suggesting its impact in SETBP1-associated disease. In addition to uncovering tissue-specific molecular signatures of *SETBP1* expression and TF activity, we provide a Shiny web application to facilitate exploring TF activity across human tissues for 758 TFs.

**Conclusions:** This study provides insight into the landscape of *SETBP1* expression and TF activity across 31 non-diseased human tissues and reveals tissue-specific expression and activity of *SETBP1* and its targets. In conjunction with the web application we constructed, our framework enables researchers to generate hypotheses related to the role tissue backgrounds play with respect to gene expression and TF activity in different disease contexts.

## Introduction

*SETBP1* is a gene located on the long (q) arm of chromosome 18 that encodes the transcription factor (TF) and oncogene SET binding protein 1 (Coccaro et al., 2017). Referred to as a DNA-binding protein, SETBP1 has several motifs, including three nuclear localization signals, a SKI homology region, and a binding region for SET nuclear oncogene. As a protein, it has a role in DNA replication and transcriptional regulation (Piazza et al., 2018). Different pathogenic variants in *SETBP1* can result in three distinct diseases (Kohyanagi & Ohama, 2023). Germline variants cause two unique ultra-rare, de novo pediatric diseases: Schinzel Giedion Syndrome (SGS) (Schinzel & Giedion, 1978) and SETBP1 haploinsufficiency disorder (*SETBP1*-HD) (Morgan et al., 2021). These conditions are differentiated by variant location, phenotypic severity, and accompanying protein gain or loss of function (GoF, LoF), respectively (Schinzel & Giedion, 1978; Morgan et al., 2021). SGS is multisystemic, involving gastrointestinal, cardiorespiratory, neurological, musculoskeletal, and urogenital abnormalities. It has a more severe phenotype than *SETBP1*-HD and affected individuals are characterized by progressive neurodegeneration and shortened life expectancy (“Schinzel Giedion Syndrome - NORD (National Organization for Rare Disorders),” 2015; Liu et al., 2018). However, SGS and *SETBP1*-HD, as disorders of protein dosage, have overlapping phenotypes, including intellectual disability, developmental delay, language impairment, distinctive craniofacial and skeletal features, and hypotonia (Schinzel & Giedion, 1978; Liu et al., 2018; Morgan et al., 2021). In contrast, somatic variants in *SETBP1* are associated with hematological malignancies and exhibit varying evidence for predisposing or promoting cancer in other adult tissue systems (reviewed in Coccaro et al., 2017).

There are multiple hypothesized mechanisms for the tissue-specificity of disease (i.e., clinical manifestations in some tissues but not others) related to intrinsic and extrinsic molecular processes spanning epigenetic, genetic, expression, regulation, and network-based mechanisms (Hekselman & Yeger-Lotem, 2020). Despite genomic advances in variant identification and sequencing technology, gaps remain in translating the role of specific genomic variants to observed phenotypic outcomes. Of these hypothesized mechanisms of tissue-specific disease manifestation, preferential or exclusive gene expression of SETBP1 targets and their altered regulation remain understudied in SETBP1-associated disorders. Potential mechanisms of SETBP1 dysfunction within neurodevelopment involving disrupted cell cycle control, DNA damage mechanisms, phosphatase activity, and chromatin remodeling have been hypothesized (Antonyan & Ernst, 2022). However, how the expression of *SETBP1* and its known TF targets function across additional tissue contexts and non-diseased human tissues requires further study.

Because of this, publicly available non-diseased data, such as from the Genotype-Tissue Expression (GTEx) project, provide an opportunity for investigating and generating hypotheses about the underlying function of disease-associated genes in different contexts, including in non-diseased and diseased contexts. Here, we investigated the gene expression of *SETBP1* and its known targets in RNA-sequencing (RNA-seq) data for each GTEx tissue. Then, we evaluated the functional enrichment of those targets based on how they clustered by expression across tissues. Next, we inferred the TF activity of SETBP1 and other TFs it is known to directly target by leveraging multivariate linear models to calculate enrichment scores representing activity for all TFs across tissues using GTEx expression and CollecTRI, a curated collection of TFs and their directional regulation on transcriptional targets (**Fig. 1**). Collectively, we have mapped the gene expression and activity of SETBP1 and its targets across human non-diseased tissues, underscoring the potential impact of tissue background. Further, we have developed a Shiny web application (https://lasseignelab.shinyapps.io/gtex_tf_activity/) to facilitate the exploration and hypothesis generation of TF activity across human tissues for 758 TFs **(Table S1)**.

**Figure 1.**
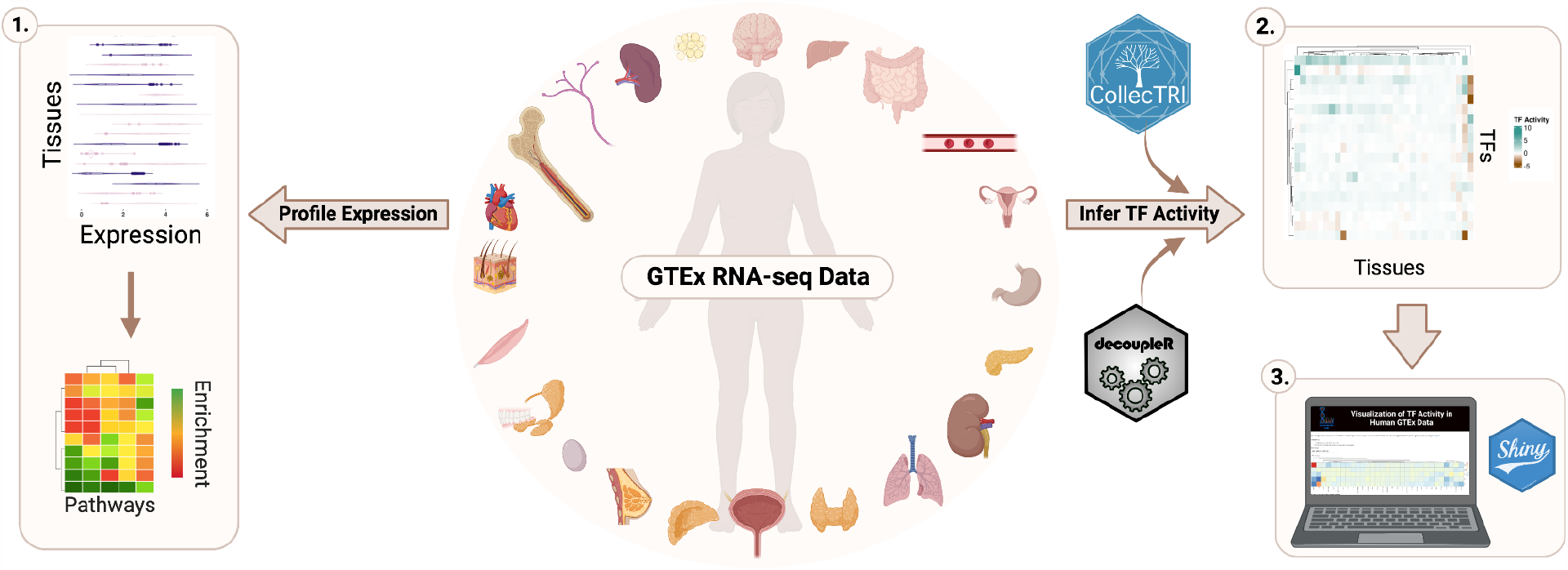
Study Overview. To investigate the tissue-specific expression and transcription factor (TF) activity of *SETBP1*, we analyzed publicly available RNA-sequencing (RNA-seq) data from the Genotype-Tissue Expression (GTEx) project. We profiled the tissue-specific expression of *SETBP1* and its known targets and performed functional enrichment analysis (1). We also inferred TF activity for SETBP1 and TFs it regulates by tissue (2) and developed an interactive web application to enable the exploration of TF activity for 758 TFs across all GTEx tissues (3).

## Materials & Methods

### Data Availability

Data and code supporting the Shiny app and to reproduce all analyses for this study are available at Zenodo (DOI:10.5281/zenodo.8222799) and https://github.com/lasseignelab/230323_JW_DiseaseNetworks (DOI:10.5281/zenodo.8225613).

The docker image used for these analyses is publicly available on Docker Hub (jordanwhitlock/setbp1_manuscript:1.0.13) and Zenodo (DOI:10.5281/zenodo.8428932). Our interactive web application can be accessed at https://lasseignelab.shinyapps.io/gtex_tf_activity/ (DOI:10.5281/zenodo.8225317).

### SETBP1 target gene set construction

We used the SETBP1 target gene set compiled in Whitlock et al. 2023 to obtain a list of known TF targets of SETBP1 (Whitlock et al., 2023). We converted a list of human HGNC to human Ensembl IDs using gprofiler2 (Kolberg et al., 2020) (v.0.2.1), resulting in a final list of 209 genes.

### RNA-sequencing expression data

Using recount3 (Wilks et al., 2021) (accessed August 2022), we obtained bulk RNA-seq data represented as transcripts per million (TPM) counts for human tissues (n=31) from the publicly available Genotype-Tissue Expression (GTEx) project. We included all tissues not classified as “study_na”.

### Classification of disease-associated affected tissues

We compiled a list of affected tissues in Schinzel Giedion Syndrome (SGS), SETBP1 haploinsufficiency disorder (SETBP1-HD), and SETBP1-associated cancer based on the literature, clinical manifestations noted within Online Mendelian Inheritance of Man (OMIM) (MIM #: 616078, 269150), and UpToDate (Schinzel & Giedion, 1978; Piazza et al., 2013; Patnaik et al., 2014; “Schinzel Giedion Syndrome - NORD (National Organization for Rare Disorders),” 2015; Elena et al., 2016; Amberger et al., 2019; Jansen et al., 2021; Morgan et al., 2021). Affected tissues for each SETBP1-associated disease phenotype included the following:

- SETBP1-HD: brain and muscle
- SGS: brain, muscle, heart, kidney, bladder, lung, small intestine, stomach, esophagus
- SETBP1-associated cancer: bone marrow and blood

While individuals with SGS and SETBP1-HD also present with vision problems and distinctive craniofacial and skeletal abnormalities, ocular and additional bone tissues are not present within GTEx v8 so they were not included in our analyses.

### SETBP1 and target gene expression and pathway enrichment

We calculated the median TPM of *SETBP1* and its targets across samples in each tissue from GTEx and scaled (log2 + 1 transformed) for visualizations. We performed complete linkage of Euclidean distances for hierarchical clustering of GTEx tissues. For target gene expression, we identified optimal k-means clusters by plotting the total within-cluster sum of squared distances between samples (inertia) for each cluster tested (k = 1-15). We identified an elbow at 3 clusters, indicating a decrease in inertia and a sufficient trade-off between information and the number of clusters captured based on the expression of *SETBP1* and targets and plotted median scaled TPM values (**Fig. S1 A**). We further verified 3 clusters to be sufficient using the Trace(W) method, which uses the trace (sum of the diagonal) of the dispersion matrix (W), finds the second differences, and selects the cluster with the maximum value between indices (**Fig. S1 B**) (Charrad et al. 2014; Milligan and Cooper 1985). To visualize this clustering and the expression of SETBP1 and its targets across GTEx tissues, we used ComplexHeatmap (version 2.10.0) (Gu, Eils & Schlesner, 2016). We next performed functional enrichment analysis (FEA) with gprofiler2 (version 0.2.1) (Reimand et al., 2007; Kolberg et al., 2020) with GO sources (GO:BP, GO:MF, and GO:CC) to identify the expression of pathways from *SETBP1* target genes for each cluster. We applied the Bonferroni procedure for multiple hypothesis correction and used a p-adjusted threshold of 0.05 and, for the background gene list, included *SETBP1* and its targets.

### TF activity

We acquired prior knowledge on the direction of TF regulation from CollecTRI (accessed May 2023) (Müller-Dott et al., 2023) and combined it with GTEx expression TPM to infer TF activity using decoupleR (Badia-I-Mompel et al., 2022) (v.2.6.0). We used a multivariate linear model (run_mlm) with a minimum threshold of 5 targets per TF to calculate activity scores (represented as t-values) for all 758 TFs. We scaled and centered data before summarizing each TF’s average regulator activity. Positive and negative scores denote TF activity and inactivity, respectively.

### Supporting Information

Table_S1.xlsx

Table_S2.xlsx

“Visualization of TF Activity in Human GTEx Data” Shiny app available at https://lasseignelab.shinyapps.io/gtex_tf_activity/

## Results

We first examined the gene expression of *SETBP1* across 31 adult human tissues from the GTEx consortium (n=19,081 samples total). We found that while *SETBP1* was expressed ubiquitously (median transcript per million (TPM) range, 0.364-16.719; **Fig. 2A**), it was most highly expressed in the cervix, blood vessel, and uterus (median TPM 16.719, 15.422, and 13.730, respectively) and most lowly expressed in the blood, bone marrow, and the adrenal gland (median TPM 0.135, 0.384, and 0.963, respectively). When we investigated the brain-region-specific expression of *SETBP1*, we found it was similarly expressed across subregions (**Fig. 2B**).

**Figure 2.**
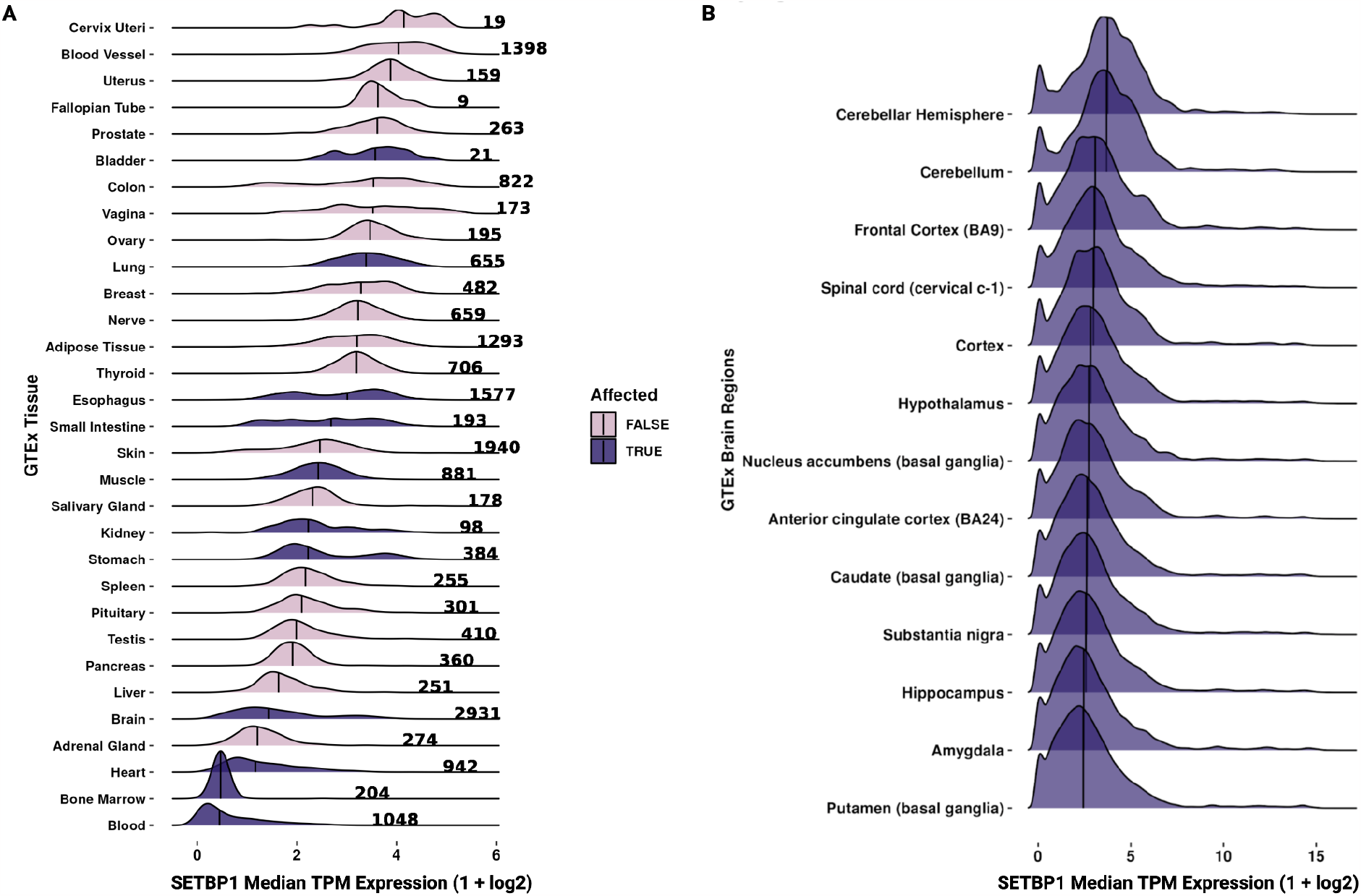
Gene expression of *SETBP1* across GTEx tissues. Ridgeline plot of the scaled median TPM values (x-axis) **(A)** across samples (denoted by bold number) for each GTEx tissue (y-axis) and **(B)** brain subregions (y-axis) for *SETBP1* where affected tissues are colored by dark purple and pink for true and false, respectively. The median is denoted by the vertical black line.

Further, we examined the gene expression of 209 known SETBP1 targets we previously compiled (Whitlock et al., 2023) from SETBP1 ChIP-seq binding sites (Kolmykov et al., 2021), MSigDB (Liberzon et al., 2015), SIGNOR (Lo Surdo et al., 2023), TRRUST (Han et al., 2018), Piazza et al. (Piazza et al., 2018), and Antonyan et al. (Antonyan & Ernst, 2022). We found that most known SETBP1 targets were also broadly expressed across adult human tissues (62.2% of known SETBP1 targets had a TPM>3 in >90% of tissues; **Fig. 3**).

**Figure 3.**
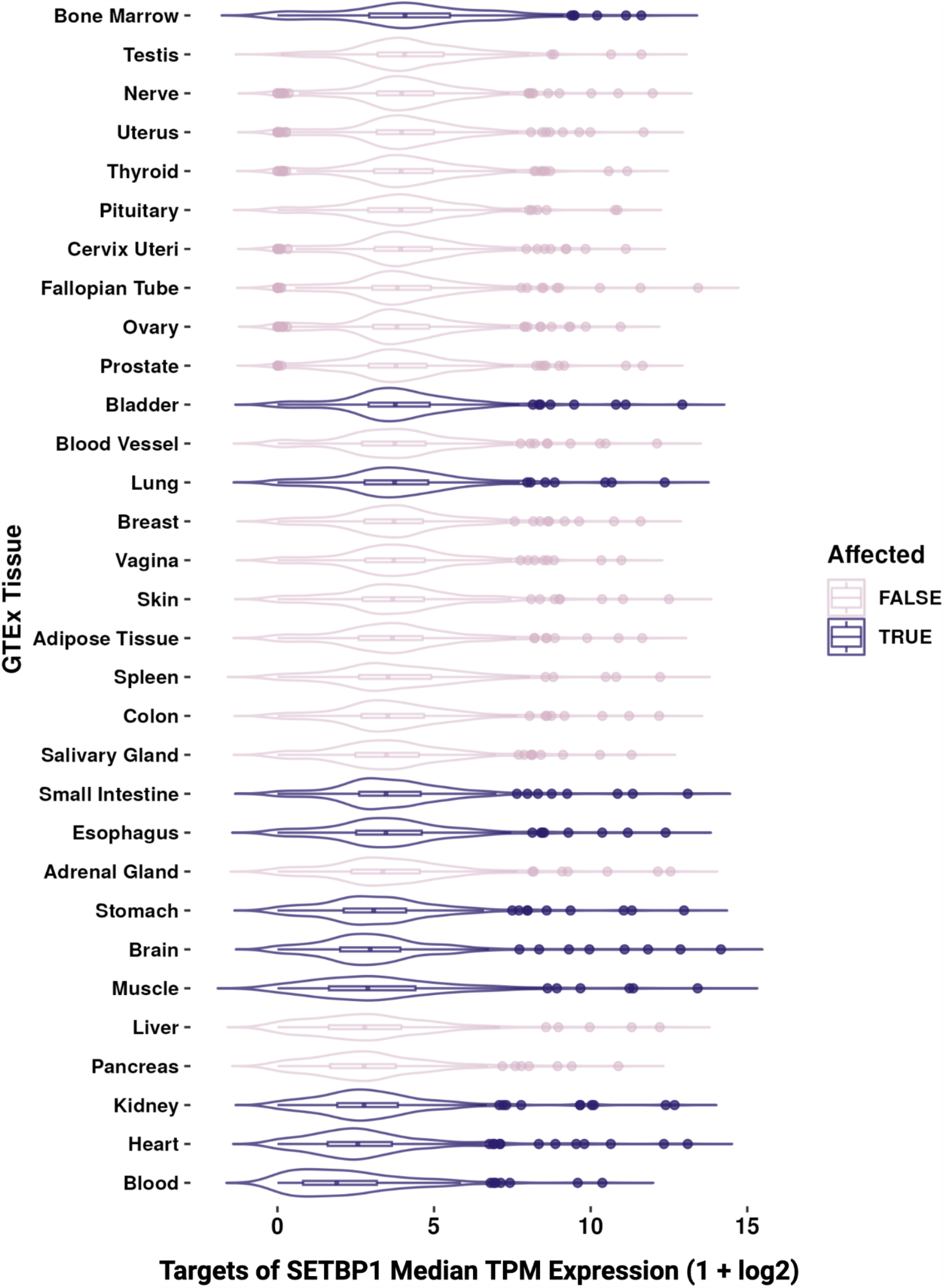
Gene expression of known SETBP1 targets across GTEx tissues. Boxplot of the scaled median TPM values (x-axis) across samples for each GTEx tissue (y-axis) for known targets of SETBP1 where affected tissues are colored by dark purple and pink for true and false, respectively.

### Targets of SETBP1 are widely expressed across tissues and cluster by gene expression into functionally distinct pathways

We next assessed the variability of SETBP1 targets’ expression across tissues and if particular functions were enriched based on their co-expression across GTEx tissues. When clustering *SETBP1* targets by gene expression, we selected three k-means clusters by both the elbow and Trace(W) method (**Fig. S1, Table S2**) and found a higher correlation of expression between *SETBP1* and its targets in clusters 1 and 2 than in cluster 3 (rho ranges -0.25 to 0.82, -0.28 to 0.84, and - 0.46 to 0.69, respectively) **(Table S2)**. The targets whose gene expression most correlated to *SETBP1* included *ZEB1* (cluster 2, rho 0.84, p-value *=* 4.63e-9), *TTC23* (cluster 2, rho 0.82, p-value *=* 1.14e-8), *RALGAPA1* (cluster 1, rho 0.82, p-value *=* 1.77e-8), *RAD52* (cluster 2, rho 0.82, p-value *=* 2.13), *CZIB* (cluster 2, rho 0.81, p-value *=* 3.07e-8), and *PBRM1* (cluster 2, rho 0.77, p-value *=* 3.54e-7). The most anti-correlated genes were the mitochondrial-associated genes *MT-RNR1* and *MT-TF* (cluster 3, rho -0.462 and -0.459, p-value *=* 0.89e-2 and 0.94e-2, respectively). Complete linkage hierarchical clustering of all tissues revealed one clade that included predominantly SETBP1-affected tissues (5 out of 7), including brain, muscle, kidney, heart, and blood (**Fig. 4A**). We further analyzed if the SETBP1 gene targets within these k-means clusters had similar functions using over-representation analysis and found that clusters 1, 2, and 3 included genes enriched for the transcription regulator complex, minor groove AT-rich DNA binding, and mitochondrial structure and function, respectively (**Fig. 4B**).

**Figure 4.**
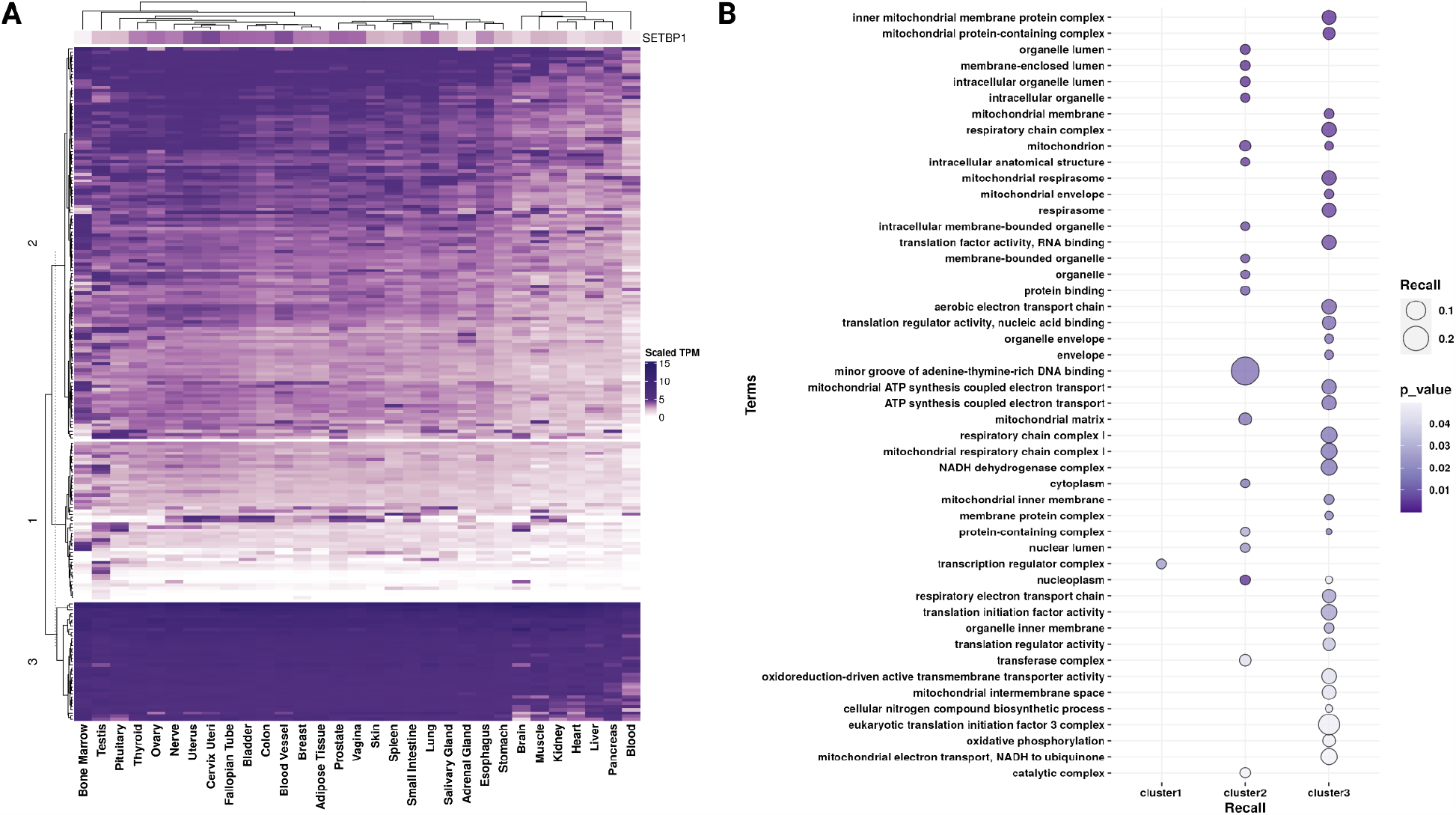
Gene expression of *SETBP1* and its known TF targets across GTEx tissues. **(A)** Heatmap clustering (y-axis) *SETBP1* and its known transcription factor (TF) targets’ scaled TPM gene expression by tissue (x-axis). **(B)** Dot plot representing the functional enrichment analysis (FEA) results for genes from clusters 1, 2, and 3 identified in A. Here dot size indicates recall, the proportion of functionally annotated input genes to each term’s full geneset size (y-axis), and color indicates significance (darker purple; more significant).

### SETBP1 TF activity is decreased in blood and increased in the pituitary

Due to its role as a TF, we next inferred the TF activity of SETBP1 across GTEx tissues. We did this by extracting the direction of TF regulation for each tissue based on prior knowledge in CollecTRI, a comprehensive, species-specific, curated database of TFs and their transcriptional targets. Using decoupleR, we then built a multivariate linear model taking into account the weight and sign of TF-target interactions and the GTEx gene expression values, where for each tissue, we interpreted positive scores as indicating SETBP1 acting as an activator, negative scores as indicating SETBP1 acting as a repressor, and zero as indicating a lack of coordinated regulation by SETBP1 **(Fig. 5)**. We found that across tissues, SETBP1 TF activity was mostly near zero (**Table S1**; range: -0.739 to 3.760 with median = -0.024 and variance = 0.53), except pituitary (TF activity = 3.760 indicating SETBP1 is functioning as an activator) and blood (TF activity = -0.739 indicating a repressive effect by SETBP1). Except for blood, we did not find that tissues known to be affected by SETBP1-associated diseases had strong evidence of SETBP1 activator or repressor roles in non-diseased adult tissues.

**Figure 5.**
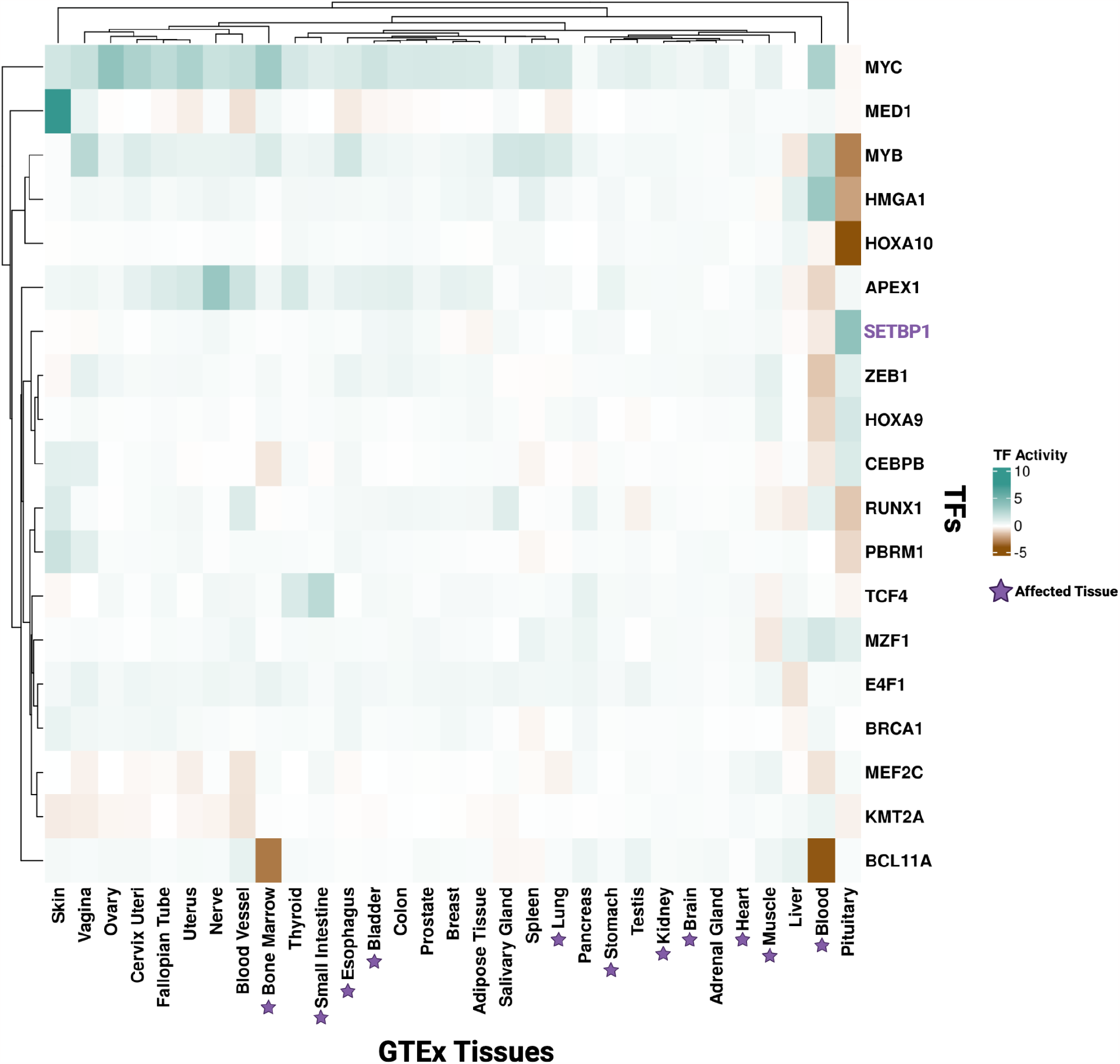
Tissue-specific TF activity of SETBP1 and TFs it targets. Heatmap representing transcription factor (TF) activity scores of SETBP1 (purple) and its known TF targets (black, y-axis) across GTEx tissues (x-axis). Teal and brown represent TF activity as an activator and repressor, respectively.

### SETBP1 TF targets demonstrate tissue-specific TF activity

In addition to SETBP1, we also inferred TF activity for targets of SETBP1 that act as TFs (**Fig. 5**). We found that these TFs, in addition to SETBP1, had a lack of coordinated TF activity in the brain, heart, and kidney, all of which are SETBP1-HD or SGS affected tissues (median = -0.004, variance = 0.008 for brain; median = -0.034, variance = 0.014 for heart; and median = -0.026, variance = 0.022 for kidney). However, SETBP1 targets associated with cancer and DNA damage had variable TF activity in the blood. For example, we predicted BCL11A (linked to multiple blood cancers as reviewed in (Bao, Cheng & Sankaran, 2019; Yin et al., 2019)) acts as a repressor in both blood and bone marrow under non-diseased conditions (TF activity -4.124 and -3.287, respectively). MEF2C (often deregulated and associated with recurrence in leukemia (Brown et al., 2018)) and ZEB1 (shown to modulate hematopoietic stem cell fates (Almotiri et al., 2021; Li et al., 2021)) also act as repressors in the blood (TF activity -0.882 and -1.532, respectively).

Furthermore, prior research showed that oncogenic apoptosis resistance and unresolved DNA damage signatures persist due to *SETBP1* variants in SGS (Banfi et al., 2021; Whitlock et al., 2023). Here, we found that in non-diseased adult tissue, TCF4, a known apoptosis regulator (Xie et al., 2012; Forrest et al., 2013), and APEX1, part of the SET complex in DNA damage response (Antonyan & Ernst, 2022), also had variable TF activity across SGS-affected tissues.

We predicted TCF4 to function as a repressor in muscle (TF activity = -0.553) and have an activator role in the small intestine (TF activity = 2.181) (**Fig. 5**). Additionally we found APEX1 had predicted repressive activity in the blood (TF activity = -1.810), but was a predicted activator in the bladder (TF activity = 0.694) and esophagus (TF activity = 0.590) (**Fig. 5**). Furthermore, we identified a lack of coordinated activity in kidney, blood, bone marrow, esophagus and muscle for TCF4 and a lack of coordinated activity in blood, muscle, and heart for APEX1 (**Fig. 5**). In summary, we have provided a map of SETBP1 and TF targets of SETBP1 activity across non-diseased adult human tissues.

## Discussion

Our study maps tissue-specific molecular signatures associated with SETBP1 across 31 human non-diseased adult tissues. We found that *SETBP1* and its known targets were widely expressed across those tissues, and FEA revealed gene sets related to transcriptional regulation, DNA binding, and mitochondrial function. Further, we uncovered tissue-specific TF activity through TF activity analysis of SETBP1 and its TF targets, underscoring the role of tissue context-driven regulation that may serve to generate hypotheses regarding the importance of TF activity in disease contexts. We also provide a framework for investigating tissue-specific gene expression and TF activity for other genes, particularly those associated with multiple diseases or multi-syndromic diseases with pathophysiological impacts across multiple tissues. As many genes associated with developmental disorders are also associated with a predisposition or increased risk of cancer (reviewed in (Nussinov, Tsai & Jang, 2022; Balachandran & Narendran, 2023)), applying this framework to those genes may be fruitful. To that end, we developed a Shiny web application with pre-computed TF activity scores that we inferred for 758 TFs for each of the 31 GTEx tissues. This application allows users to search for TFs and compare their activity across all 31 GTEx tissues (https://lasseignelab.shinyapps.io/gtex_tf_activity/).

With complete-linkage hierarchical clustering of SETBP1 target gene expression by tissue, we discovered one clade largely consisted of affected tissues (5 out of 7 of the tissues in the clade), including brain, kidney, heart, and blood, the most frequently noted tissues impacted by SETBP1 perturbations (**Fig. 4A**) (Schinzel & Giedion, 1978; Matsumoto et al., 2005; Acuna-Hidalgo et al., 2017; Coccaro et al., 2017). These results suggest that, in non-diseased tissues, there is a similarity in the expression of SETBP1 targets across multiple tissues known to exhibit a phenotype in SETBP1-associated diseases. We further investigated the trends of expression of SETBP1 targets by k-means clustering and the correlation of each target’s expression to *SETBP1* gene expression (**Fig. 4A, Table S2**). We compared the total within-cluster sum of squares (“elbow” method) as well as Trace(W) method to select 3 k-means clusters (Charrad et al. 2014; Milligan and Cooper 1985). We highlighted the highest correlated and anti-correlated genes associated with SETBP1’s roles as an activator or a repressor (Coccaro et al., 2017). Of note, some of the most significantly correlated and anti-correlated targets included genes with known critical roles in development (*ZEB1*, p-value *=* 4.63e-9(Drápela et al., 2020) and *RALGAPA1*, p-value *=* 1.77e-8 (Wagner et al., 2020)), with additional involvement in metastasis and therapy resistance for *ZEB1* (Drápela et al., 2020). Among the non-significant highest correlated and anti-correlated genes were *HMGA1* (p-value *=* 0.13) and *MYB* (p-value *=* 0.17). Their non-significance in correlation analyses to SETBP1 across tissues may highlight a potential context dependence or tissue-specific regulatory roles. In addition to having functions in organism development, HMGA1 enhances recovery from double-stranded DNA breaks. When overexpressed, it sensitizes cells to DNA damage and is a driver of malignant tumors (Fujikane et al. 2016). Additionally, SETBP1’s consensus binding site largely overlaps with the AT-hook consensus motif of HMGA1 (Piazza et al. 2018). On the other hand, *MYB* expression is known to be critical for myeloid leukemia induced by SETBP1 activation, and its inhibition could be beneficial for treating SETBP1-associated neoplasms (Coccaro et al. 2017; Nguyen et al. 2016). The genes we identified here highlight previously known mechanisms underlying SETBP1-associated disease and provide additional potential targets for future investigation.

To uncover potential functional patterns by expression clusters, we also tested each of our 3 k-means clusters of gene targets for over-representation of GO terms where recall represented the proportion of functionally annotated input genes to each term’s full gene set size (**Fig. 4B**). We found cluster 3 genes were ubiquitously and highly expressed across tissues. These genes were enriched for terms regarding mitochondrial structure and function (**Fig. 4B**) and included many mitochondrially-encoded *SETBP1* targets (*MT-ND6, MT-RNR1, MT-TE, MT-TP, MTCO3P12, MT-TF*) (**Table S2**). Cluster 2 exhibited the highest recall with GO terms enriched for minor groove AT-rich DNA binding (**Fig. 4B**), similar to regulation by SETBP1 as well (Piazza et al., 2018). Cluster 1 genes enrichment was for just one GO term, transcription regulator complex (**Fig. 4B**), and included genes for the cancer-associated transcription factors *RUNX1, HOXA9*, and *MYB* (Bhagwat & Vakoc, 2015). These results further highlight the varying roles of SETBP1 targets and their expression patterns across tissues.

Out of all SETBP1 disease-associated tissues, blood was the only tissue exhibiting notable SETBP1 TF repression. Previous studies support that surpassing a higher functional threshold (i.e., more damaging or impacting variants) is required for SETBP1-driven cancer (Acuna-Hidalgo et al., 2017). Our findings here with respect to SETBP1 in non-diseased blood, suggest that GoF variants (already known to drive blood-associated cancers such as myeloid leukemia (Albano et al., 2012; Piazza et al., 2013; Acuna-Hidalgo et al., 2017)) may impact SETBP1’s TF repressive role, suggesting future research directions that may shed light on the mechanism behind SETBP1-associated blood cancers. For example, SETBP1 has been shown to activate *MYC (Carratt et al., 2022)*, which we calculated has a TF activity score of 2.77 in blood (activator role) and positive TF activity scores in 27 out of 30 other tissues (**Fig. 5**). Likewise, we and others (Banfi et al., 2021; Antonyan & Ernst, 2022; Whitlock et al., 2023) have hypothesized that in the presence of pathogenic germline variants, alterations in TCF4 or APEX1 activity lead to apoptosis resistance or increased DNA damage in SGS. Our results suggest both exhibit activator roles across many tissues in non-diseased adult tissues, so disease-associated perturbations may impact SETBP1 TF activity, contributing to SGS.

Our study relies on the assumption that TF protein activity can be inferred by the weighted mode of regulation and transcript levels of its target genes (Badia-I-Mompel et al. 2022). A major limitation of this study is that we conducted analyses in adult bulk expression profiles. For disease-associated genes with developmental phenotypes like *SETBP1*, temporal expression (Cardoso-Moreira et al., 2019, 2020) and TF activity are likely key to linking disrupted genes to molecular and physiological phenotypes. Comprehensive prenatal and postnatal gene expression atlases in active development (e.g., developmental GTEx, dGTEx) will provide an unprecedented opportunity to repeat and expand the analyses in this study across developmental time points. Additionally, as we conducted the analyses here in bulk profiles, we cannot assess the gene expression and TF activity of particular cell types. However, as we recently reported, SETBP1’s role as an epigenetic hub leads to cell-type-specific differences in TF activity, gene targeting, and regulatory rewiring in the mouse cerebral cortex and kidney (Whitlock et al., 2023). This underscores the importance of future studies that generate and analyze the necessary data to understand cell-type-specific gene expression and TF activity across human tissues. Furthermore, some affected tissues related to the skeleton and eyes are not included within GTEx v8. If this data becomes available, future studies could investigate tissue-specific expression and activity for vision loss, craniofacial, and skeletal abnormalities within SGS and SETBP1-HD. Finally, the GTEx tissues may have subclinical pathologies that have not been previously reported, so care must be taken with the interpretation of non-affected tissues.

## Conclusions

In summary, our study highlights the importance of considering tissue-specific expression and regulatory properties in investigating disease-related genes. It provides a basis for future investigations of TFs involved in processes across many tissues, including developmental and cancer contexts.

## List of Abbreviations

FEA: Functional Enrichment Analysis
GTEx: Genotype-tissue expression
GoF: gain of function
LoF: loss of function
OMIM: Online Mendelian Inheritance of Man
RNA-seq: RNA-sequencing
SETBP1-HD: SETBP1 haploinsufficiency disorder
SGS: Schinzel Giedion Syndrome
TF: Transcription Factor
TPM: Transcripts per Million

## Acknowledgements

The authors thank the Lasseigne Lab members Vishal Oza, Tabea Soelter, Emma Jones, and Victoria Flanary for their feedback throughout this study. We also thank the UAB Biological Data Science group (RRID:SCR_021766) for providing a script for helping to run containers on the UAB high-performance cluster (https://github.com/U-BDS/training_guides/blob/main/run_rstudio_singularity.sh)

## Authors’ contributions

Conceptualization: JHW, EJW, TCH, ADC, and BNL. Methodology: JHW and BNL. Software: JHW (including the Shiny app) and EJW. Validation: JHW, EJW, TCH, and ADC. Formal Analysis: JHW and EJW. Investigation: JHW, EJW. Resources: BNL. Data Curation: JHW. Writing - Original Draft: JHW. Writing - Review & Editing: JHW, EJW, TCH, ADC, and BNL. Visualization: JHW and EJW. Supervision: BNL. Project Administration: BNL. Funding Acquisition: BNL.

## Ethics approval and consent to participate

Not applicable.

## Consent for publication

Not applicable.

