## Supplementary material for "The landscape of *SETBP1* gene expression and transcription factor activity across human tissues": Supp Figure 1

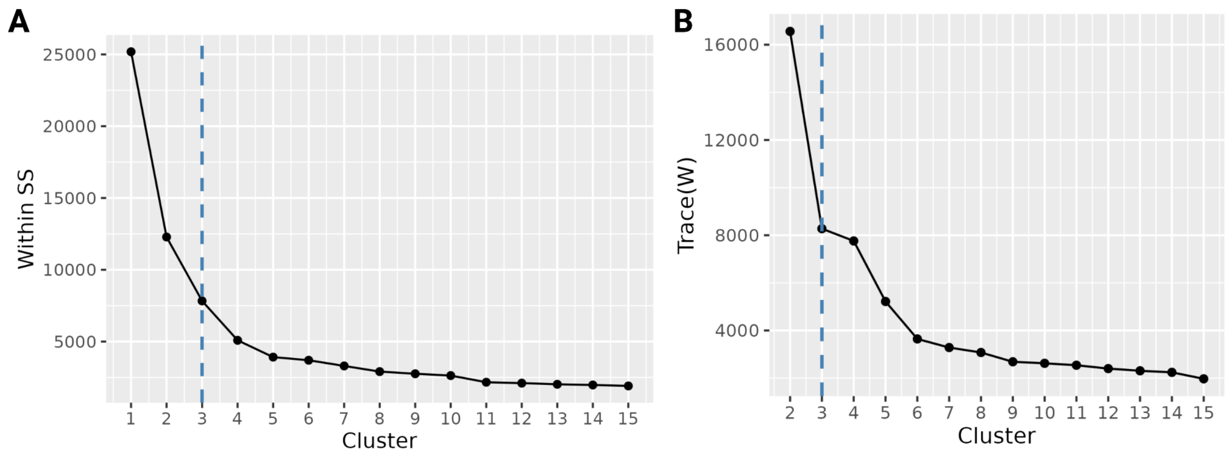


**Fig S1. Determining optimal k-means clusters**. K-means clustering indices of GTEx scaled normalized *SETBP1* and gene targets' expression using (**A**) Elbow plot, 1-15 k-means clusters (x-axis) plotted by their total within-cluster sum of squared distances (inertia), where dashed blue line signifies the point at which the inertia decreases and represents a sufficient number of clusters. (**B**) Line plot of Trace(W), the sum of the diagonal of the sum of squared within-group dispersion matrix (y-axis) for each cluster (x-axis) is used to calculate second differences, and the optimal cluster (dashed blue line) is indicated as the maximum value between levels.
